# STAR-Fusion: Fast and Accurate Fusion Transcript Detection from RNA-Seq

**DOI:** 10.1101/120295

**Authors:** Brian J. Haas, Alex Dobin, Nicolas Stransky, Bo Li, Xiao Yang, Timothy Tickle, Asma Bankapur, Carrie Ganote, Thomas G. Doak, Nathalie Pochet, Jing Sun, Catherine J. Wu, Thomas R. Gingeras, Aviv Regev

## Abstract

**Motivation:** Fusion genes created by genomic rearrangements can be potent drivers of tumorigenesis. However, accurate identification of functionally fusion genes from genomic sequencing requires whole genome sequencing, since exonic sequencing alone is often insufficient. Transcriptome sequencing provides a direct, highly effective alternative for capturing molecular evidence of expressed fusions in the precision medicine pipeline, but current methods tend to be inefficient or insufficiently accurate, lacking in sensitivity or predicting large numbers of false positives. Here, we describe STAR-Fusion, a method that is both fast and accurate in identifying fusion transcripts from RNA-Seq data.

**Results:** We benchmarked STAR-Fusion’s fusion detection accuracy using both simulated and genuine Illumina paired-end RNA-Seq data, and show that it has superior performance compared to popular alternative fusion detection methods.

**Availability and implementation:** STAR-Fusion is implemented in Perl, freely available as open source software at http://star-fusion.github.io, and supported on Linux.

## Introduction

Major efforts are underway to dissect the molecular underpinnings of cancer development, progression, evolution, and resistance to therapies (Cancer Genome Atlas Research, et al., 2013; International Cancer Genome, et al., 2010). Research has shown there are many paths to tumorigenesis, often involving genetic mutations leading to activation of oncogenes and inactivation of tumor suppressors (Hanahan and Weinberg, 2000; Hanahan and Weinberg, 2011). Chromosomal rearrangements leading to the formation of fusion transcripts represent one class of genomic aberrations that occur at high frequencies in certain cancer types, including leukemias and prostate cancer (Yoshihara, et al., 2015), as well as produce strong genetic drivers of the disease across a wide range of cancers (Stransky, et al. 2014). Among the known cancer-driving fusion transcripts with high prevalence in specific cancer types are BCR—ABL1, found in ~95% of chronic mylogenous leukemia patient samples (Lim, et al., 2005), TMPRSS2—ERG in ~50% of prostate cancers (Tomlins, et al., 2005), and DNAJB1—PRKACA found to be the hallmark and likely driver of fibrolamellar carcinoma, a rare liver tumor affecting young individuals (Honeyman, et al., 2014). Determining the driver of a given tumor is important to inform the best therapeutic strategy and match drugs effective against certain oncogenic fusions with the patients who could benefit from them. For example, tyrosine kinase inhibitors have been highly effective in the treatment of tumors harboring kinase fusion transcripts in leukemia and other cancers (Zhao, et al., 2015) (Shaw and Solomon, 2015) (Druker, et al., 2006) (Gross, et al., 2015).

Genome and transcriptome sequencing are powerful methods to capture evidence for chromosomal rearrangements and fusion transcripts. Unlike point mutations and indels, which can be readily captured from efficient Whole Exome Sequencing (WES), genome sequencing evidence for fusion genes typically requires Whole Genome Sequencing (WGS). Conversely, transcriptome sequencing by RNA-Seq is highly suited for fusion transcript detection, since it represents the “expressed exome” of the tumor itself: it captures only the transcriptionally active regions of the genome, reflects any rearrangements, and reduces sequencing costs compared to WGS, thus focusing fusion-finding efforts on actively transcribed gene fusions that are most likely relevant to tumorigenesis.

Several bioinformatics methods and software tools have been developed to identify candidate fusion transcripts from RNA-Seq (reviewed in (Latysheva and Babu, 2016; Wang, et al., 2013)). The general strategies fall into two conceptual classes: mapping-first approaches that align RNA-Seq reads to genes and genomes to identify discordantly mapping reads suggestive of rearrangements, and assembly-first approaches that directly assemble reads into longer transcript sequences followed by identifying chimeric transcripts consistent with chromosomal rearrangements. Evidence supporting predicted fusions is typically measured by the number of RNA-Seq fragments found as split (junction) reads that directly overlap the fusion transcript chimeric junction, or as spanning fragments (bridging read pairs) where each paired read maps to opposite sides of the chimeric junction without directly overlapping the chimeric junction itself (shown in Figure 1).

**Figure 1:**
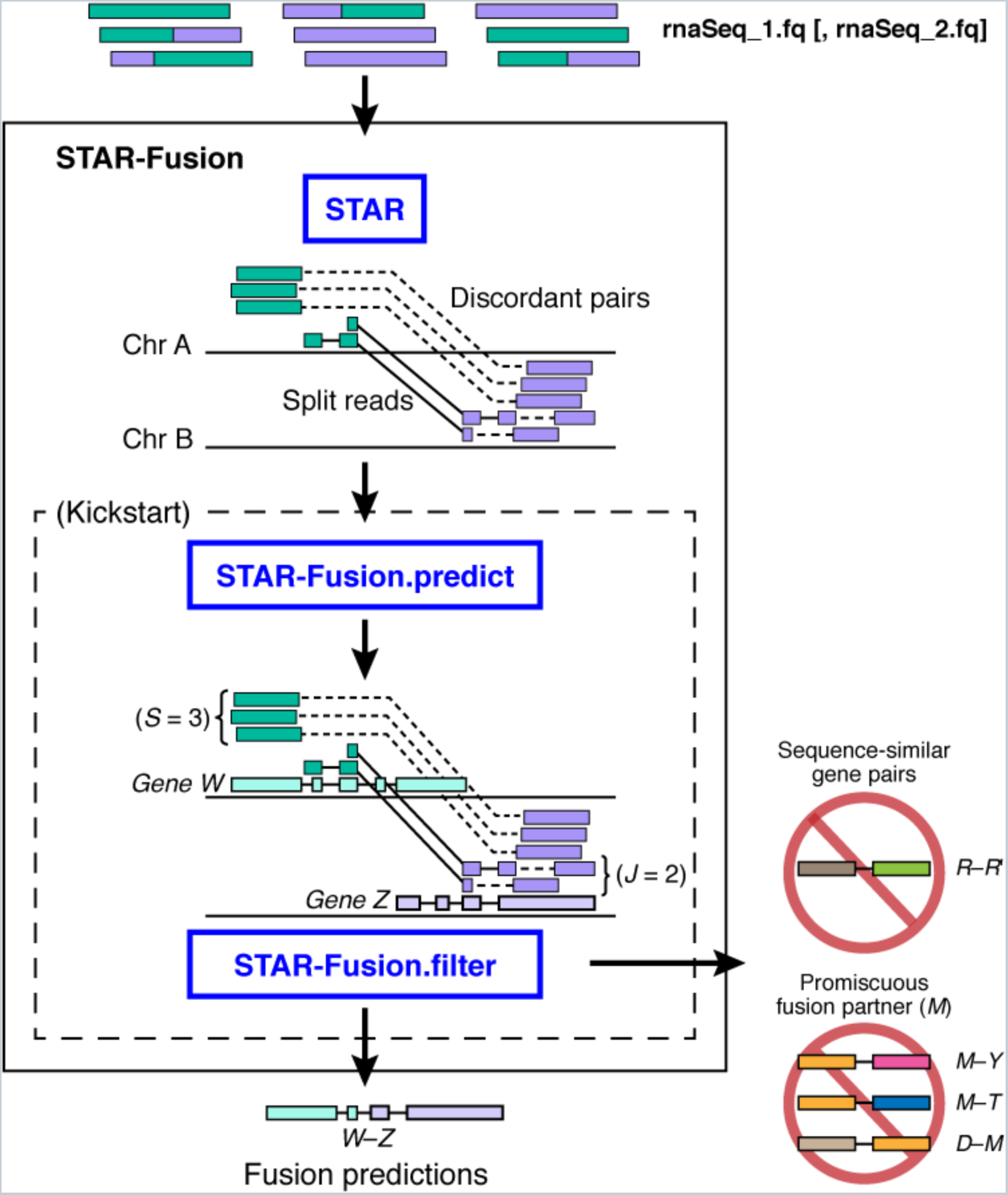
Overview of the STAR-Fusion pipeline. Illumina RNA-Seq reads are aligned to the genome using STAR. Discordant and split-read alignments are identified, mapped to reference transcript structure annotations, filtered to remove likely artifacts, and scored according to the abundance of fusion-supporting reads. Fusion candidates containing sequence-similar gene pairs or promiscuous fusion partners are excluded as likely false positives.

Implementations of the various prediction methods vary in the read alignment tools, the genome database and gene set resources used, criteria for reporting candidate fusion transcripts and excluding false positives, all resulting in varying prediction accuracies in addition to installation complexity, execution time, robustness, and hardware requirements. Depending on the fusion prediction tool chosen, processing a single RNA-Seq sample containing tens of millions of reads can take several days worth of computing and result with a list of hundreds to thousands of gene fusion candidates, most of which are likely false, having little evidence supporting their prediction. Faster and more accurate methods for fusion detection are urgently needed, particularly as RNA-Seq ventures into the realm of precision medicine and clinical care.

A critical component of mapping-first approaches is the read alignment step. In particular, the STAR aligner was originally developed as a fast and accurate RNA-Seq alignment tool with capabilities to report split reads and discordantly aligned pairs (Dobin, et al., 2013; Dobin and Gingeras, 2015; Dobin and Gingeras, 2016). Based on these capabilities, STAR was incorporated into a fusion detection pipeline to explore the landscape of kinase fusion transcripts in cancer (Stransky, et al., 2014), demonstrating that STAR could be leveraged to identify novel driver fusion transcripts. STAR has since been incorporated into ensemble fusion prediction pipelines (Nicorici, et al., 2014).

Here, we describe STAR-Fusion – a fusion prediction tool leveraging the speed and accuracy of the STAR RNA-Seq aligner. STAR-Fusion builds upon and significantly expands a previously described successful strategy for calling fusions (Stransky et al, 2014), and is implemented here as a freely available open source software tool. STAR-Fusion performs a fast mapping of fusion evidence to reference transcript structure annotations and filters likely artifacts to report accurate fusion predictions. We demonstrate that STAR-Fusion provides accurate fusion transcript prediction using both simulated and genuine RNA-Seq data derived from cancer cell lines, and show that it is both more accurate and substantially faster than other alternatives.

## Methods

### STAR-Fusion pipeline

The STAR-Fusion pipeline (Figure 1) takes Illumina RNA-Seq data (ideally paired-end, but compatible with single-end data) as input and generates lists of candidate fusion transcripts as output. It also requires several genomic resources which are packaged into a Genome Resource Lib: the reference genome, reference transcript structure annotations, and results from an all-*vs*-all BLAST+ (Camacho, et al., 2009) search of reference transcript sequences. The Genome Resource Lib is compiled once for a given set of genomic resources and simply pointed to as a parameter to STAR-Fusion at run time. Genome Resource Libs are provided for human and mouse, and instructions and software are available to facilitate compiling a new Genome Resource Lib for a different reference genome or different reference transcript annotation set (below). We discuss each of the components in turn.

### Aligning RNA-Seq reads to the genome using STAR, capturing split and discordantly mapping reads

The chimeric detection in STAR is implemented as an extension of the STAR standard mapping procedure. In the first step, the maximum mappable prefix algorithm is used to find the seeds exactly matching to the genome. Next, genomic alignment windows are selected by clustering the anchor seeds, which are defined as those mapping to the genome fewer times than a user-defined parameter (--*winAnchorMultimapNmax=50* by default). The maximum window size determines the maximum genomic length of the local alignment, and is controlled by *--alignMatesGapMax* and *--alignIntronMax*. In each genomic window, a local alignment of the read sequence is performed by stitching together all seeds within the window. If the best alignment among all windows does not cover the entire read sequence, chimeric detection is performed by finding the next best scoring window that covers the remainder of the read sequence. Thus, the resulting chimeric alignment consists of two “segments”, each being a local linear alignment, with the minimum lengths restricted by *--chimSegmentMin* to prevent output of unreliable chimeras with short alignments. If a chimeric segment contains an intervening splice junction proximal to the chimeric junction, the minimum sequence segment length for the chimeric junction-proximal segment is specified by *--chimJunctionOverhangMin*. The maximum unmapped read sequence in-between two chimeric segments is limited by *--chimSegmentReadGapMax*.

The STAR-Fusion mapping parameters are based upon the best practices for STAR (Dobin and Gingeras, 2015; Dobin and Gingeras, 2016), as well as parameters optimized to capture fusion genes, as described above (Stransky, et al., 2014):

> “STAR --chimSegmentMin 12 --chimJunctionOverhangMin 12 --chimSegmentReadGapMax 3 --alignSJDBoverhangMin 10 --alignMatesGapMax 200000 --alignIntronMax 200000 --alignSJstitchMismatchNmax 5 -1 5 5 --twopassMode Basic”

### Mapping split and discordant read alignments to reference gene annotations (STAR-Fusion.predict)

To determine the candidate gene pairs of potential fusions, the discordant read pairs and split read alignments reported by STAR are next mapped to exons of reference transcript annotations based on coordinate overlaps, leveraging interval tree data structures (as implemented in the Perl CPAN module Set::IntervalTree (http://search.cpan.org/~benbooth/Set-IntervalTree-0.01/lib/Set/IntervalTree.pm) for fast coordinate overlap queries. STAR-Fusion selects those candidate gene pairs for which the fusion-supporting evidence indicates a sense-sense orientation between the fusion pairs and scores them according to the number of split reads supporting the fusion breakpoint and the number of paired-end fragments that span the breakpoint. Duplicate paired-end read alignments are removed.

While most filtering of fusion candidates is performed by the STAR-Fusion.filter step as described below, some initial filtering based on read support is performed here simply to remove the weakest supported candidates and limit the subsequent more rigorous STAR-Fusion.filter step to a smaller number of candidates. This improves overall computing efficiency and reduces unnecessary input/output. STAR-Fusion applies minimal read support criteria of at least one junction-defining split read and at least two total supporting fragments (sum of junction reads and spanning fragments). If the chimeric junction occurs at a position that does not coincide with a reference annotated exon splice junction, then at least three breakpoint reads are required. All minimal support criteria are configurable as program parameters.

### Filtering fusion predictions (STAR-Fusion.filter)

Finally, STAR-Fusion determines the most likely correct fusions, filtering out unlikely candidates from the initial predictions. To this end, the candidates are further grouped according to gene pairings and breakpoint proximity and then further filtered according to read support, extent of alignment at the putative breakpoint, and sequence similarity between putative fusion gene partners, performed in the following order:

1. **Grouping by breakpoint proximity**. Fusions with non-reference breakpoints that fall within a window of +/-5 bases of a dominantly supported breakpoint are merged into single fusion predictions with the dominant breakpoint.
2. **Assessing strength of alignment evidence**. Those fusions that have only split-read support (lacking spanning fragment support) are required to have at least 25 bases of read alignment at each end of the breakpoint, though not necessarily from a single read, as different reads can provide the required alignment overlap at either end.
3. **Filtering lowly supported fusion isoforms**. If multiple fusion isoforms are predicted for a given gene pair, we discard those isoforms having less than 10% of split-read supporting evidence compared to the dominantly supported breakpoint.
4. **Filtering sequence-similar fusion pairs**. We exclude fusion candidates A—B where there exist a significant sequence alignment (BLAST, E<=10^−3^) between any two reference isoforms of the genes. Instead of performing BLAST at runtime, the results from an all-*vs*-all BLAST search are stored and indexed for immediate retrieval at runtime.
5. **Filtering promiscuous fusion partners**. Finally, promiscuous fusion genes, defined as those gene partners found to have multiple fusion partners predicted within a single sample, are filtered in two stages: first, those fusions containing a fusion gene that has less than 20% supporting evidence than a dominant fusion pair containing that same gene are discarded; second, any fusion pair containing a partner observed in at least 3 remaining fusion predictions are altogether excluded.

Those fusions that pass the above filters are reported in a tab-delimited summary file identifying the fusion pairs, the inferred fusion breakpoint (chromosomal exon boundaries), counts of supporting split reads and spanning fragments, and identification of the RNA-Seq reads that support the fusion prediction.

### Compiling a Genome Resource Lib

Compiling a Genome Resource Lib for use with STAR-Fusion requires three inputs: (1) a reference genome in FASTA format, (2) reference transcript structure annotations in GTF format, and (3) results from an all-*vs*-all BLAST search of repeat-masked transcript sequences. Note that repeat masking is required in order to avoid detecting sequence similarities between transcripts that derive from mobile elements and other repeats that often occur within noncoding regions such as within lncRNAs or untranslated regions of otherwise unrelated coding transcripts. The results from the all-*vs*-all BLAST search are stored into a Berkeley DB for fast querying based on gene identifiers. We provide full instructions and helper utilities for building the Genome Resource Lib at http://fusionfilter.github.io.

In the work described here, the Genome Resource Lib was constructed with the reference genome and transcript annotations in Gencode Release 19 (https://www.gencodegenes.org/releases/19.html), and restricted to the reference chromosomes (1-22, X, Y, and MT). Reference transcript sequences were reconstructed based on the GTF annotation file and then repeat masked using RepeatMasker (Smit A.F.A) as follows:

> “RepeatMasker -pa 6 -s -species human -xsmall ref_annot.cdna”

The repeat masked transcript sequences were then searched all-*vs*-all for sequence similarity using BLAST as follows:

> “blastn -query ref_annot.cdna.masked -db ref_annot.cdna.masked -max_target_seqs 10000 -outfmt 6 - evalue 1e-3 -lcase_masking -num_threads 20 -word_size 11 > blast_pairs.outfmt6”

The genome was then indexed by STAR as follows:

> “STAR --runThreadN 4 --runMode genomeGenerate --genomeDir ref_genome.fa.star.idx --genomeFastaFiles ref_genome.fa --limitGenomeGenerateRAM 40419136213 --genomeChrBinNbits 16 --sjdbGTFfile ref_annot.gtf --sjdbOverhang 100”.

### Benchmarking Fusion Prediction Accuracy

We assessed fusion prediction accuracy using simulated and genuine RNA-Seq data, and compared it to 15 previously developed methods. Specifically, we downloaded and installed each of PRADA (Torres-Garcia, et al., 2014), TopHat-Fusion (Kim, et al., 2013; Kim and Salzberg, 2011), deFuse (McPherson, et al., 2011), nFuse (McPherson, et al., 2012), EricScript (Benelli, et al., 2012), ChimeraScan (Iyer, et al., 2011), FusionCatcher (Nicorici, et al., 2014), FusionHunter (Li, et al., 2011), JAFFA-Assembly, JAFFA-Hybrid, and Jaffa-Direct (Davidson, et al., 2015), SOAPfuse (Jia, et al., 2013), MapSplice (Wang, et al., 2010), ChimPipe (Rodriguez-Martin, et al., 2017), and InFusion (Okonechnikov, et al., 2016). To ensure consistency, we reconfigured SOAPfuse and TopHat-Fusion to leverage the Gencode v19 annotation. All fusion prediction methods with the exception of JAFFA-Assembly leverage read alignment to target reference genomes or transcriptomes, whereas JAFFA-Assembly first performs *de novo* assembly to reconstruct transcripts and then aligns the reconstructed transcripts to target sequences to identify likely fusions. JAFFA-Hybrid leverages both direct read mapping and *de novo* transcriptome assembly (Davidson, et al., 2015). Programs and parameters used are provided in **Supplementary Table 1**. Benchmarking data, scripts, and the analysis protocols followed are provided at https://github.com/STAR-Fusion/STAR-Fusion_benchmarking_data and in the **Supplementary Text**.

### Simulated fusion transcripts and RNA-Seq data

We generated simulated chimeric transcripts using custom scripts, developed and released here as the Fusion Simulator Toolkit (https://FusionSimulatorToolkit.github.io). FusionSimulator selects two protein-coding genes at random from the Gencode v19 annotations. It then constructs a fusion transcript by randomly fusing a pair of exons randomly selected from each gene, requiring that each gene contributes at least 100 bases of transcript sequence to the generated fusion, and that the fusion breakpoint occurs between two exons that have consensus dinucleotide splice sites. In generating a set of fusion genes, any gene participating as a fusion partner is allowed to exist in only one fusion pair.

We simulated RNA-Seq reads using ‘rsem-simulate-reads’ in the RSEM software (Li and Dewey, 2011). RSEM was first used to estimate the expression values of the Gencode v19 reference transcripts supplemented with the simulated fusion transcripts. Next, the expression values of the simulated fusion transcripts were reset randomly according to a log2 distribution of TPM expression values in the dynamic range of 1 to 15. Simulated read lengths and read quality characteristics were modeled based on genuine RNA-Seq data sets as described below. After directly setting fusion transcript expression values, all transcript expression values were renormalized to TPM values (summing to 1 million) and subject to RNA-Seq read simulation using ‘rsem-simulate-reads’.

This process was applied separately for ten samples, each generating 500 random fusions and simulating 30 million PE Illumina RNA-Seq reads. Half of the simulated samples generated 50 base reads and the other half 101 base reads, the former modeled on short RNA-Seq reads generated by the Illumina Human Body Map 2.0 study (ArrayExpress study E-MTAB-513; http://www.ebi.ac.uk/arrayexpress), and the latter based on a set of cancer cell lines from the Cancer Cell Line Encyclopedia (CCLE) (Barretina, et al., 2012) (sources for the targeted data sets are listed in **Supplementary Table 2**). Simulated fusion transcripts and simulated RNA-Seq data are made available at https://data.broadinstitute.org/Trinity/STAR_FUSION_PAPER/SupplementaryData/sim_reads/.

### Fusion prediction in cancer cell line transcriptomes

Paired-end Illumina RNA-Seq data were obtained from 60 publicly available cancer cell line data sets, spanning a variety of cancer types (data sources and representative cancer types are listed in **Supplementary Table 3**). Cancer cell lines included 52 from the CCLE project, and further supplemented with 8 other cancer cell lines popularly studied for fusion detection including: the breast cancer cell lines BT474, KPL4, MCF7, and SKBR3 (Edgren, et al., 2011); VCaP (prostate cancer); LC2/ad and H2228 (lung adenocarcinoma); and K562 (erythroleukemia). To facilitate benchmarking and runtime analysis, 20 million paired-end reads were randomly sampled from each data set and targeted for fusion prediction. All sampled cancer cell line RNA-Seq data targeted for fusion discovery are available at https://data.broadinstitute.org/Trinity/STAR_FUSION_PAPER/SupplementaryData/cancer_cell_lines/FASTQ/.

### Fusion prediction accuracy computation

True positive (TP), false positive (FP), and false negative (FN) fusion predictions were assessed for each method. The true positive rate (TPR; or recall or sensitivity), positive predictive value (PPV, precision), and F1 accuracy measure (the harmonic mean of TPR and PPV) were computed according to standards:

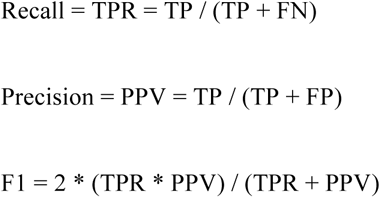

TP and FP were assessed at each minimum supporting evidence threshold to generate precision-recall curves, and prediction accuracy was measured as the area under the precision-recall curve (AUC), which is better suited than the popular receiver operating characteristic curve for studies such as fusion prediction where the numbers of true negatives (at least ~20k^2^, considering possible gene pairings) far exceeds the number of true positive fusions (Davis, 2006).

Additional details regarding benchmarking methods, software, result files, and links to input RNA-Seq data and related resources are provided in the **Supplementary Text** and at https://github.com/STAR-Fusion/STAR-Fusion_benchmarking_data.

## Results

### STAR-Fusion accurately and sensitively predicts fusion on simulated data

We compared STAR-Fusion and the 15 other methods on ten sets of simulated RNA-Seq data sets, each containing 30 M PE reads and each including 500 fusion transcripts expressed at a broad range of expression levels. To examine the effect of read length on fusion prediction accuracy, half the samples contained 50 base reads and the other half contained 101 base reads, reflective of read lengths of contemporary RNA-Seq data sets.

Most methods performed well on the simulated RNA-Seq data with high recall (TPR) and precision (PPV), with top contenders having slight differences in prediction accuracy (Figure 2a, **Supplementary Tables 4-7**). Most predictors had improved performance with longer (101 base) reads. Notably, JAFFA-Direct and JAFFA-Hybrid were restricted for use with longer reads (Davidson, et al., 2015). Most fusions that were not captured had the lowest expression levels and hence had little to no corresponding read evidence.

**Figure 2.**
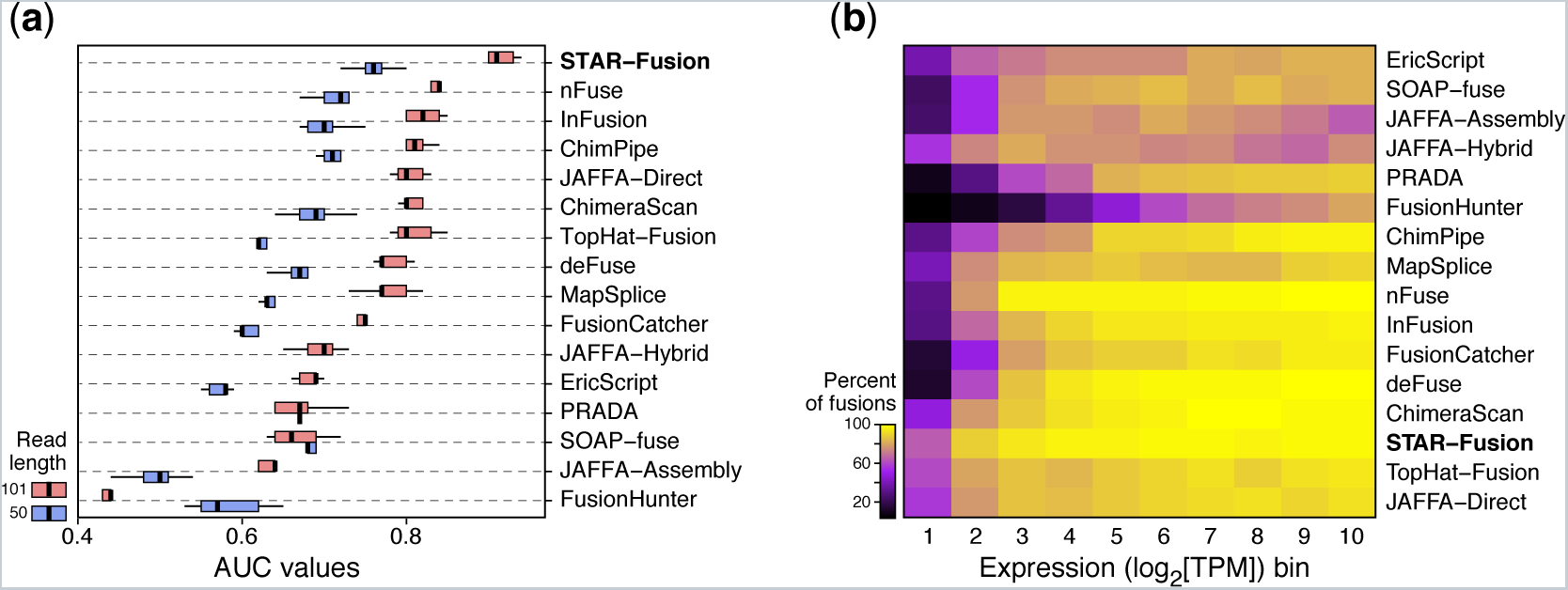
Benchmarking of fusion prediction accuracy on simulated data. A. Distribution of precision-recall AUC values for each predictor using short (blue, 50 base) or longer (red, 101 base) simulated reads. Boxplots show the distribution of AUC values among the 5 replicates, with the median drawn as the center line, boxes extending to the 1st and 3rd interquartile, and whiskers extending to furthest data points within 1.5 * the interquartile range. B. Sensitivity of fusion prediction (recall) for simulated fusions at different expression levels. Shows is the percent of simulated fusions (color bar) identified by each method (rows) at each expression level (log_2_(TPM)) bin, where bins range from 2^1^ to 2^10^. Ordering of rows in the heatmap is based on hierarchical clustering.

STAR-Fusion had superior accuracy with high sensitivity for fusion detection across the full dynamic range of gene expression (Figure 2, **Supplementary Tables 4-8**). Since *de novo* assembly generally requires higher sequence coverage for transcript reconstruction and fusion detection, JAFFA-Assembly had limited sensitivity for fusion prediction. However, it is peculiar that JAFFA-Assembly demonstrated reduced sensitivity at the highest fusion transcript expression levels; this may result from JAFFA-Assembly using the Oases *de novo* transcriptome assembler, as similar approaches leveraging Trinity for *de novo* assembly continued to retain high sensitivity for the same highly expressed fusion transcripts (data not shown), and the assembly-free JAFFA-Direct did not exhibit this aberrant behavior.

The vast majority of false positives obtained on the simulated data were due to paralogous genes of fusion transcripts being reported in addition to or in place of the known fusion gene; eg. Fusion *A—B* reported as *A’—B* where *A’* is a paralog of gene *A*. Scoring fusions containing paralogs as true positives improves reported prediction accuracies for all methods (Supplementary Fig 1), with STAR-Fusion continuing to demonstrate top performance.

### STAR-Fusion accurately and sensitively predicts fusions on cancer cell line transcriptomes

Next, we compared the fusion transcript predictions of STAR-Fusion and each of the 15 methods in 60 cancer cell line transcriptomes. Several of the 60 cell lines are well studied and include fusion transcripts that have been well documented and experimentally validated (Edgren, et al., 2011) (Suzuki, et al., 2013). Nevertheless, unlike with simulated data, there is no perfect truth set by which prediction accuracies can be computed with absolute certainty. Instead, we used concordance between methods as a proxy for accuracy: we hypothesized that those fusions predicted by multiple different predictors are the most likely to be true and those uniquely predicted by a given predictor are more likely to be false. Based on this concordance benchmarking approach, we defined the truth set as those fusions agreed upon by at least four different methods, and those uniquely predicted by a single method as false positives. Those predictions agreed upon by only 2 or 3 methods were treated as ambiguous (neither TP nor FP).

Unlike simulated data, there was much more substantial variation in the performance of different methods on the cancer cell line data, particularly in the numbers of putative FPs observed across methods (Figure 3, **Supplementary Tables 9, 10**). The numbers of uniquely predicted fusions ranged from the tens to over a thousand at points of maximum prediction accuracy (peak F1 score in Figure 3D and **Supplementary Table 9**). Not all of the methods that performed well on simulated data were observed to excel on the cancer cell line data; for example, nFuse was among the top performers using simulated data (Figure 2), but lacked predictive power on the cancer cell line data (Figure 3).

**Figure 3.**
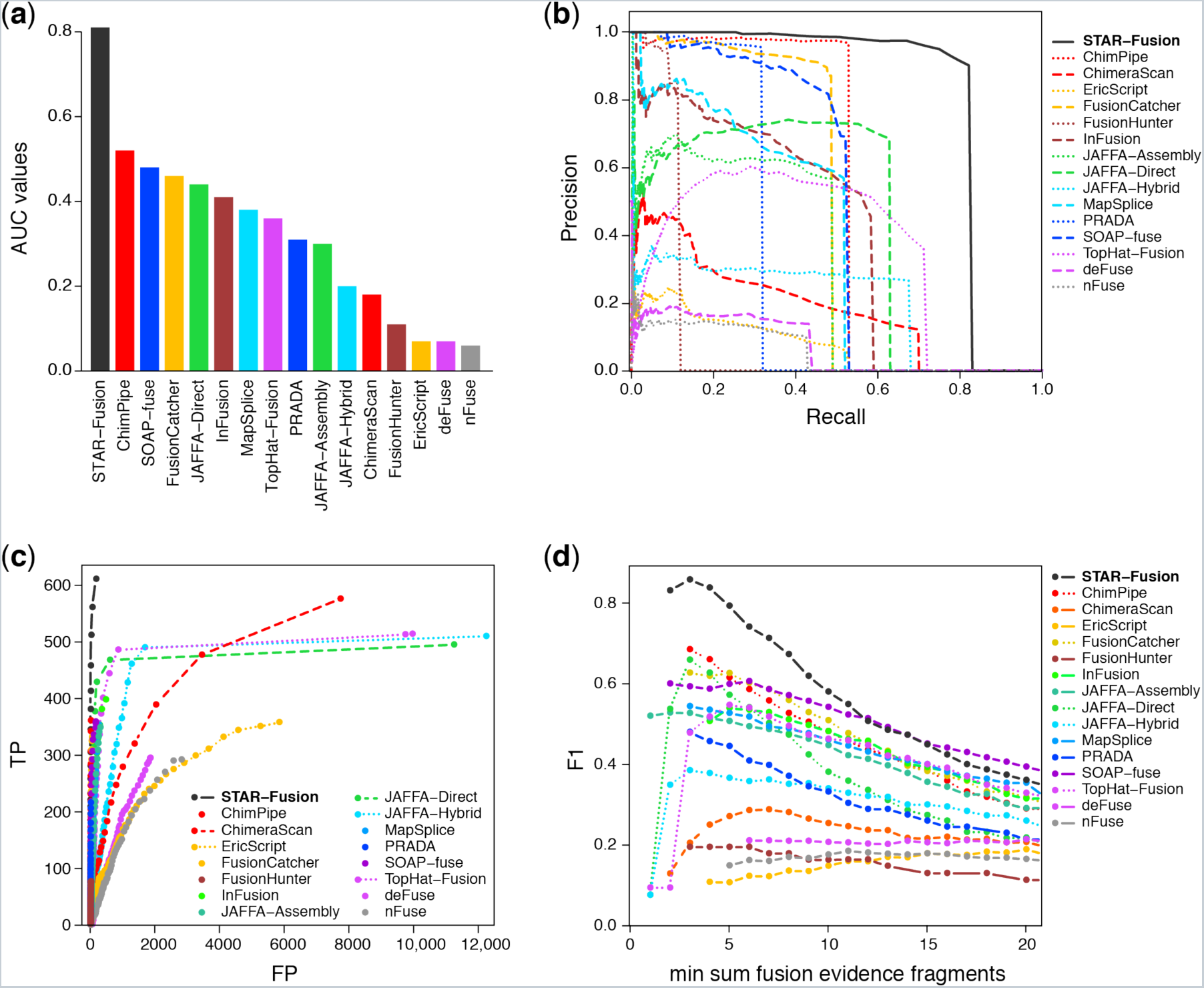
Benchmarking of fusion prediction accuracy on cancer cell line transcriptomes. A. Precision-recall curve AUC values for each method ranked from highest to lowest. B. Precision-recall curves from which AUC values were computed. C. True positives (TP, Y axis) and false positives (FP, X axis) at each minimum fusion-supporting read evidence thresholds. D. F1 accuracy measurements (Y axis) at each minimum read evidence thresholds (X axis).

Nevertheless, STAR-Fusion was the top performing method on these data, having the highest AUC, followed by the recently published ChimPipe software. Alternative definitions of truth sets might change the relative rankings of methods, but STAR-Fusion continues to demonstrate superior performance (Supplementary Figure 3 and **Supplementary Text**). For example, defining the truth set as fusions agreed upon anywhere between 2 and 5 methods, ignoring ambiguous predictions or treating ambiguous predictions as FP, and/or considering paralogs as proxies to TP predictions had no effect on STAR-Fusion’s top ranking among all methods.

Fusion prediction accuracy highly depends on the minimum threshold of evidence for split reads and junction-bridging fragments. For example, STAR-Fusion has peak accuracy on the cancer cell line data at a minimum of three total fusion-supporting RNA-Seq fragments (Figure 3D), whereas TopHat-Fusion reaches peak accuracy at a minimum of 5 fragments, but exhibits a steep decline in accuracy when lowering the threshold to 3 fragments (Figure 3D), due to an influx of false positives with few additional true positives (Figure 3C). Notably, STAR-Fusion exhibited better performance across a range of minimum evidence thresholds (Figure 3D). In general, requiring at least 3 supporting RNA-Seq fragments per 20 M total reads (or normalized to 0.15 fusion fragments per million total RNA-Seq fragments (FFPM)) appears to provide near-optimal fusion prediction accuracy for the well-behaved prediction methods in this study.

### STAR-Fusion prediction pipeline is fast

Finally, we compared the run time of each fusion prediction pipeline on conventional hardware leveraging the computing grid at the Broad Institute, with each of the 60 cancer cell line samples run as a separate compute job for each fusion prediction tool (Figure 4). Run times for fusion predictors on single samples (20 M PE reads) ranged from hours to days.

**Figure 4.**
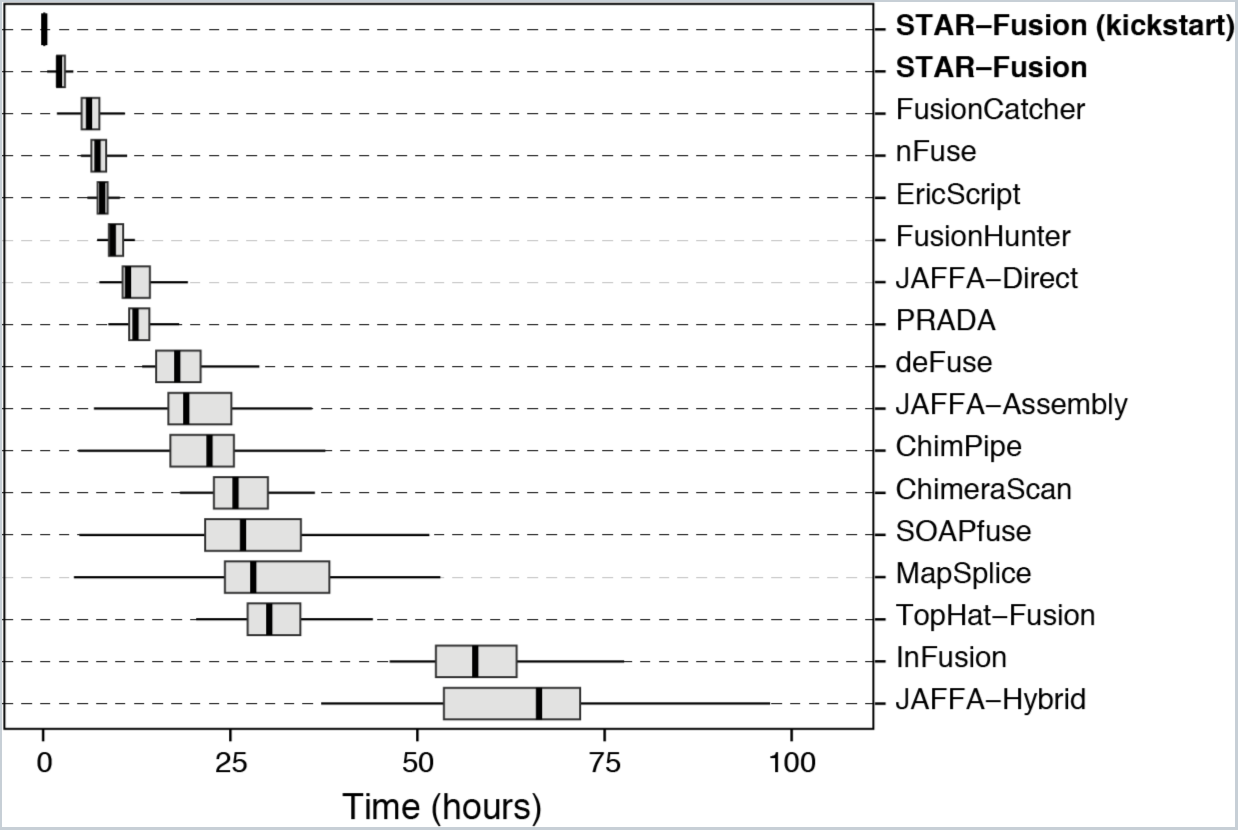
Distribution of run times for fusion predictors on the 60 cancer cell line data sets. Boxplots are drawn to represent the median, interquartile range, and whiskers drawn to furthest data points within 1.5 * the interquartile range.

STAR-Fusion has the shortest run time, taking on average 2.3 hours per sample, given a single compute core; STAR-Fusion can run at approximately 30 minutes per sample when provided with 10 compute cores (Supplementary Figure 4). Most of STAR-Fusion’s execution time involves running STAR to align the RNA-Seq reads to the reference genome. Mapping the split read alignments and discordantly aligned pairs to the reference annotations and subsequent filtering of fusion candidates is comparably fast, taking only 9 minutes per sample on average.

## Discussion

Fast and accurate fusion detection is essential to ensure progress in cancer research and to bring next-generation sequencing to the precision medicine pipeline. The National Cancer Institute Genomic Data Commons (https://gdc-portal.nci.nih.gov), current home for The Cancer Genome Atlas data, now contains RNA-Seq samples for over 10,000 tumors and totaling over 100 TB of data, and with current trends, the data deluge is expected to grow exponentially. Further cancer genome sequencing is expected as it becomes more commonly deployed in the clinical setting and routinely used for treatment decision-making and prognostics (Gagan and Van Allen, 2015; Horak, et al., 2016). We hope that the advances enabled by STAR-Fusion will enable it to play an important role in identifying known and actionable fusion transcripts in the clinical context, in addition to discovering new hallmarks of diverse cancer types in pan-cancer research studies.

Fusion transcript discovery is just one aspect of the application of next-gen transcriptome sequencing to cancer research. RNA-Seq and genome read alignment already exist as standard methods in cancer sequencing studies. Due to the fast speed and accuracy of the STAR aligner (Dobin, et al., 2013; Engstrom, et al., 2013), STAR is already being used as a key component of production pipelines for aligning reads to genomes and estimating gene expression (https://www.encodeproject.org/pipelines/ENCPL002LPE/), and STAR is an integral component of the GATK best practices for variant calling using RNA-Seq data (https://software.broadinstitute.org/gatk/guide/article?id=3891). By enabling the STAR option to report the chimeric read alignments, these data will be readily available for running STAR-Fusion in the faster ‘kickstart’ mode, and quickly provide access to lists of candidate fusion transcripts.

STAR-Fusion is freely available as open source and included with documentation and sample data at http://star-fusion.github.io. We also provide access to running STAR-Fusion on user-provided RNA-Seq data via a point-and-click interface at our Trinity Cancer Transcriptome Analysis Toolkit Galaxy Portal: https://galaxy.ncgas-trinity.indiana.edu.

## Acknowledgements

We thank Daniel Nicorici, author of FusionCatcher, for inspiration and guidance. We thank Leslie Gaffney for assistance with graphical illustrations.

---

**Supplementary Figure 1:**
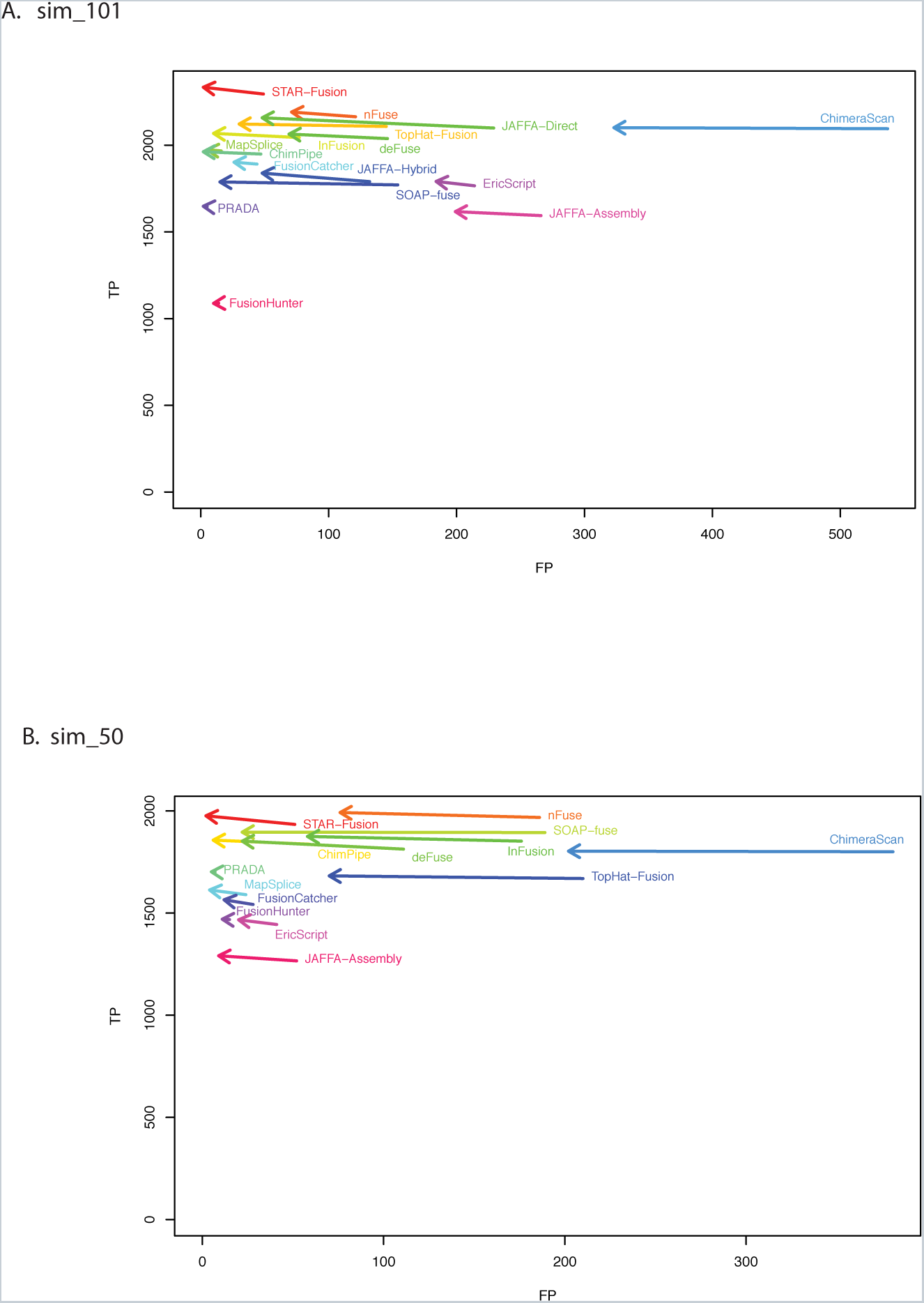
Fusion prediction TP and FP before and after treating paralogs of true fusion partners as equivalent. The impact is most notably a decrease in the number of false positives with a small increase in the number of true positives. Results are shown for the **A**. simulated 101 base length PE reads and **B**. simulated 50 base length PE reads.

**Supplementary Figure 2:**
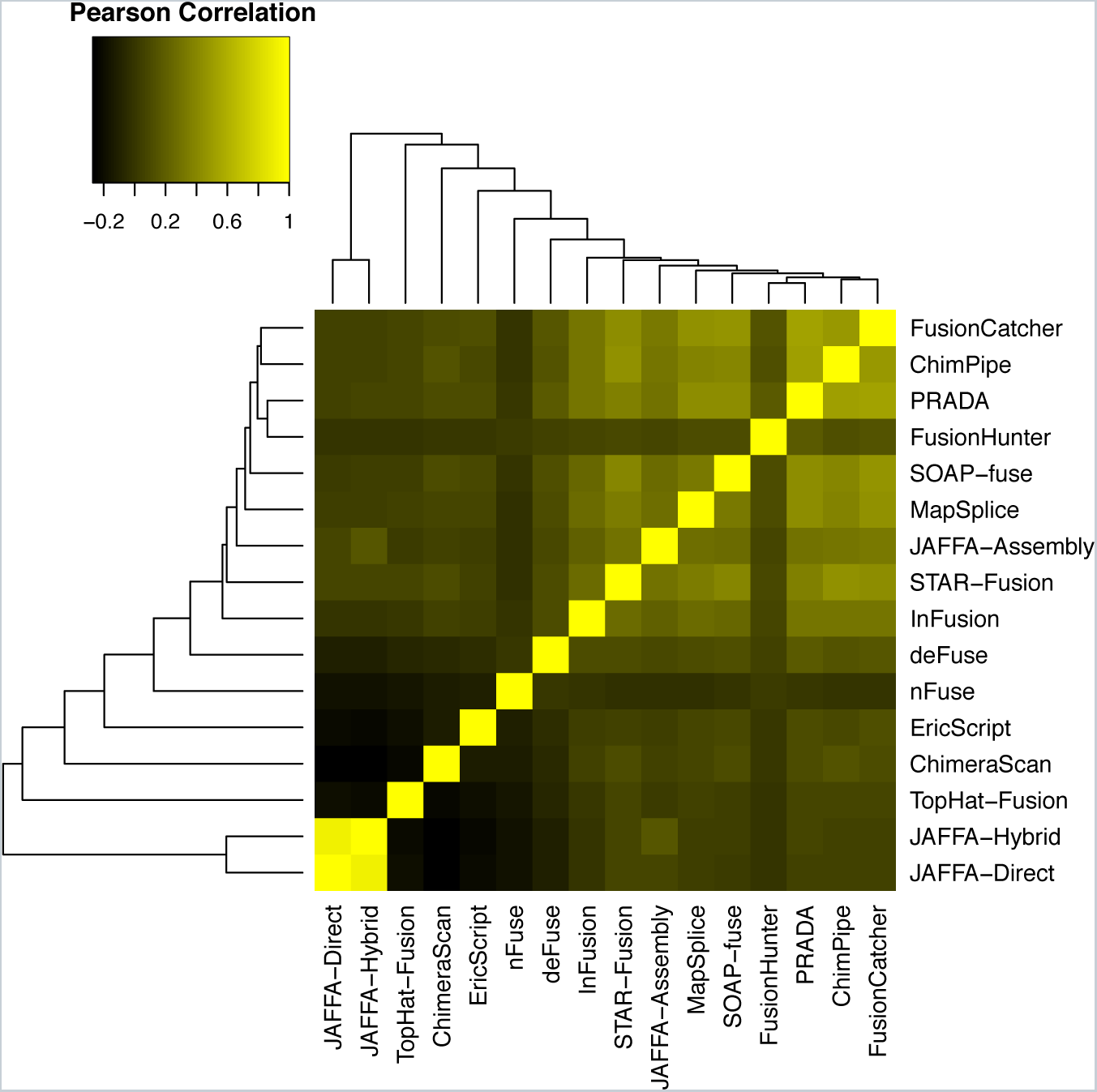
Correlation of predicted fusion gene pairs among different fusion prediction methods for the 60 cumulative cancer cell line data sets. Only JAFFA-Direct and JAFFA-Hybrid demonstrate high correlation among predictions, and so JAFFA-Hybrid was excluded from the majority voting process of defining candidate truth sets to avoid unintended bias.

**Supplementary Figure 3:**
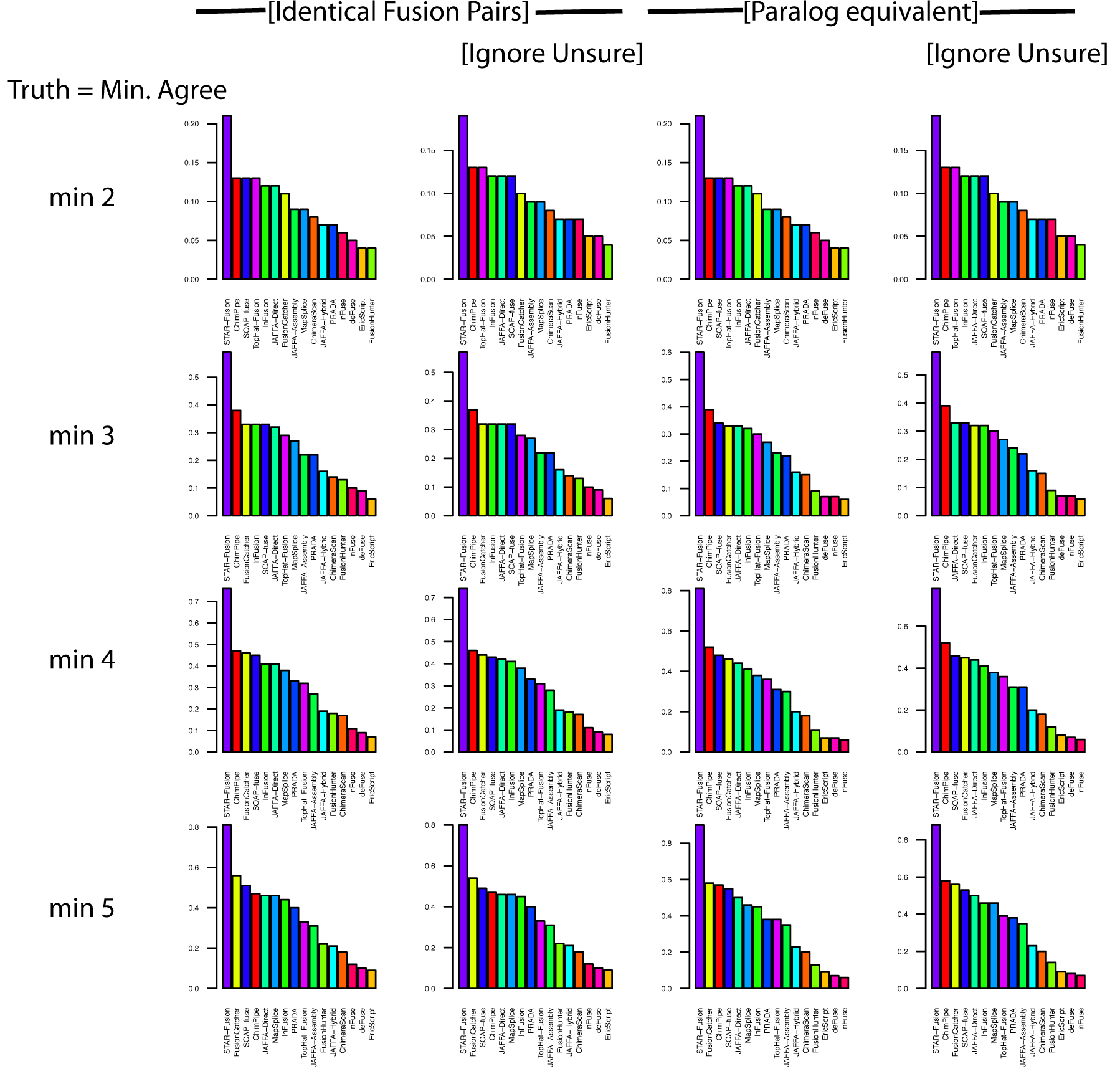
Fusion prediction accuracy (AUC) computed using different truth set definitions. The minimum threshold for the number of methods that must agree to define the truth set is shown at left and ranges from 2 to 5. True positive fusions are identified by strict gene identity (A--B) (left two columns) or allowing paralogs of fusion partners to suffice as proxies to the true fusion partner (A’--B’ where A’ is either A or a paralog of A, and B’ is either B or a paralog of B). In all cases, uniquely predicted fusions are treated as false positives. In the columns indicated ‘Ignore Unsure’, those predictions agreed upon by >1 predictor and < threshold predictors are ignored and not counted as TP, FP, or FN.

**Supplementary Figure 4:**
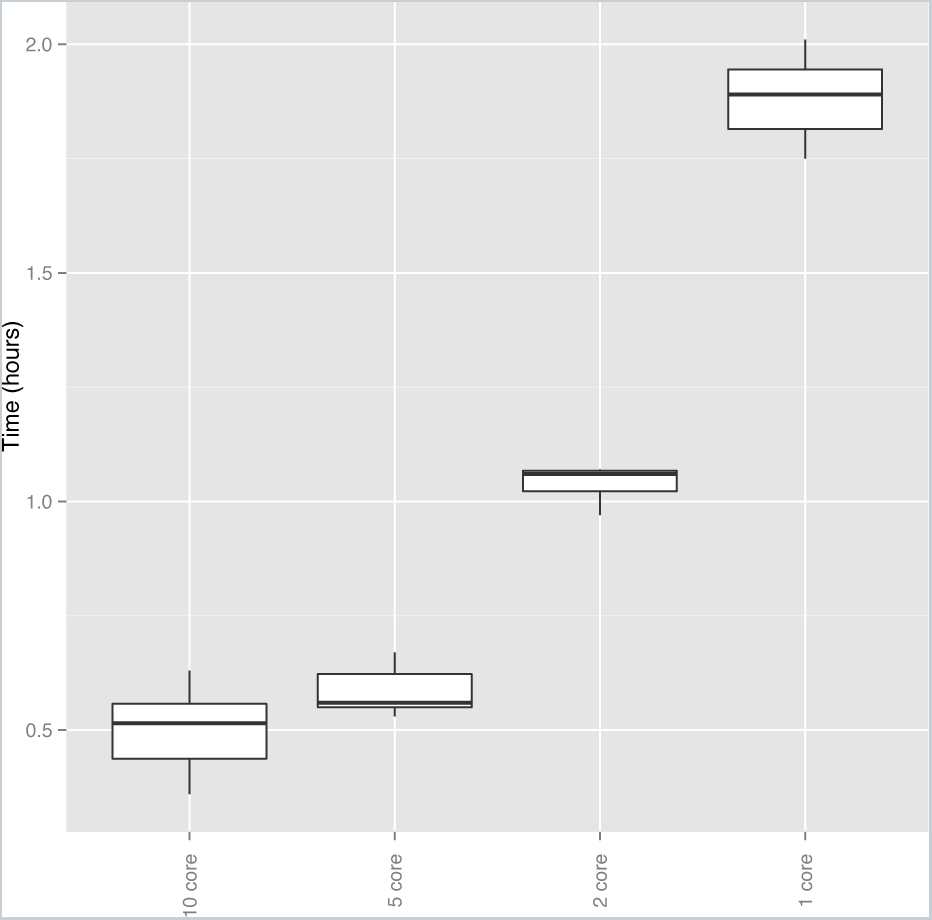
STAR-Fusion runtime using multiple threads. The STAR aligner was run as part of STAR-Fusion with multithreading set to the specified number of threads and total STAR-Fusion execution time was recorded. Using 10 threads, runtime was approximately 30 minutes.

